# Combined sensor-based monitoring of mycothiol redox potential and DNA-damage response in *Corynebacterium glutamicum*

**DOI:** 10.1101/2022.07.25.501298

**Authors:** Fabian Stefan Franz Hartmann, Ioannis Anastasiou, Tamara Weiß, Tsenguunmaa Lkhaasuren, Gerd Michael Seibold

## Abstract

Excessive amounts of reactive oxygen species (ROS) can cause irreversible damages to essential cellular components such as DNA. Genetically encoded biosensors targeting oxidative stress and DNA-stress have emerged to a powerful analytical tool to assess physiological states in a non-invasive manner. In this study, we aimed to combine the redox biosensor protein Mrx1-roGFP2 with a transcriptional biosensor for DNA-damage based on the P*_recA_* promoter fused to a reporter gene (*e2-crimson*) in *Corynebacterium glutamicum*. Therefore, the redox biosensor strains *C. glutamicum* WT_Mrx1-roGFP2 and the mycothiol (MSH)-deficient mutant strain *C. glutamicum* Δ*mshC*_Mrx1-roGFP2 were equipped with the DNA-stress reporter plasmid pJC1_P*_recA__e2-crimson*. Exposure of the double-sensor equipped *C. glutamicum* WT strain to hypochlorite resulted in an oxidative redox shift, accompanied by an induction of the DNA-stress reporter system. In absence of the major non-enzymatic antioxidant MSH, the induction of the DNA-stress response was even more pronounced. This confirms the linkage of oxidative stress and DNA-damage response, and therefore making antioxidants a crucial player to protect DNA. Furthermore, exposure of the double biosensor strains to a DNA-damage inducing agent resulted in an oxidative redox shift. These results suggest a direct link of DNA-damage and oxidative stress response in *C. glutamicum*. Finally, we observed that inhibition of cell wall biosynthesis by penicillin caused both an oxidative redox shift and a DNA-damage response in *C. glutamicum*. The excellent compatibility of Mrx1-roGFP2 with E2-Crimson shown here provides a powerful combinatorial biosensor concept for in-depth studies of redox-related physiology in future studies.

## 1 Introduction

The Gram-positive *Corynebacterium glutamicum* is widely used in industrial biotechnology for amino acid production and has been engineered into a versatile platform organism for the production of bulk and fine chemicals (Eggeling and Bott, 2015; Wolf *et al*., 2021). During aerobic fermentation processes, *C. glutamicum* unavoidably becomes exposed to conditions leading to an accumulation of reactive oxygen species (ROS). Excessive amounts of ROS can cause irreversible damages to essential cell components such as proteins, lipids or nucleic acids (Antelmann and Helmann, 2011; Imlay, 2013; Ezraty *et al*., 2017). To counteract oxidative stress, *C. glutamicum* is equipped with effective enzymatic and non-enzymatic ROS detoxification systems (Chi *et al*., 2014; Su *et al*., 2018; Liu *et al*., 2021). The non-enzymatic antioxidant mycothiol (MSH), for instance, has been shown to be essential to maintain a reducing environment in the cytoplasm of *C. glutamicum* (Van Loi *et al*., 2015; Tung *et al*., 2019). Accordingly, inactivation of genes essential for MSH biosynthesis has detrimental effects on the organisms’ performance when facing oxidative stress conditions (Hartmann *et al*., 2020).

In addition, bacteria often initiate DNA repair systems upon ROS exposure as shown for *Escherichia coli* (Goerlich *et al*., 1989; Halliwell *et al*., 2021), *Caulobacter crescentus* (Leaden *et al*., 2018) and *Acinetobacter baumanii* (Aranda *et al*., 2011). Consequently, mutant strains lacking DNA-repair systems are often more susceptical towards oxidative stress (Imlay and Linn, 1987; Ajiboye *et al*., 2018). In general, DNA-damage leads to the occurrence of single-stranded DNA (ssDNA), which is recognized by the protein RecA. Upon interaction with ssDNA, RecA gets activated via oligomerization which allows the self-cleavage of the transcriptional repressor LexA. This, in turn, promotes the expression of DNA repair genes (Montelone, 2006; Aertsen and Michiels, 2006; Erill *et al*., 2007; Kreuzer, 2013). The linkage of oxidative stress and DNA-damage response is known as oxidative DNA-damage (Poetsch, 2020; Halliwell *et al*., 2021).

Adequately assessing the “stress-status” of microbial cells is a difficult task to undertake for in-depth physiological studies. Current research studies often make use of laborious omics technologies to capture changes of the transcriptome, proteome or metabolome upon applying artificial stressors (Chi *et al*., 2014; Pedre *et al*., 2015; Van Der Heijden *et al*., 2016; Liu *et al*., 2016). While such omics technologies provide comprehensive insights on a global scale, it is not well suited towards gathering time-resolved data at a cellular level. Genetically encoded biosensors have emerged to a powerful analytical tool to easily assess time-resolved physiological states in a non-invasive manner (Polizzi and Kontoravdi, 2015; Carpenter *et al*., 2018). For instance, the redox active versions of GFP (roGFP1 and roGFP2) allow to detect changes of the spectral characteristics depending on the oxidation state of the fluorescent protein (Dooley *et al*., 2004). Redox active proteins harbor two Cys residues that form a disulfide bond upon oxidation, resulting in ratiometric changes of two excitation maxima in the fluorescence excitation spectrum. These properties make them well suited as ratiometric oxidative stress biosensors (Schwarzländer *et al*., 2016). Recently, roGFP1 and roGFP2 were fused *via* a polypeptide linker to functionally active domains involved in redox sensing such as Glutaredoxin-1 (Grx1) or Mycoredoxin-1 (Mrx1). Such biosensor variants allow to selectively equilibrate with low-molecular-weight (LMW) thiols such as mycothiol (2MSH/MSSM; Mrx1-roGFP2) or glutathione (2GSH/GSSG; Grx1-roGFP2) (Bhaskar *et al*., 2014; Schwarzländer *et al*., 2016). As LMWs are the major redox buffer systems utilized by organisms, their quantitative assessment provides a good indicator with respect to the overall redox environment in a cell. Reporter systems based on components of the SOS response are commonly utilized for the detection of DNA-damage response in bacterial cells (Vollmer *et al*., 1997; Van Der Meer and Belkin, 2010). For studies in *C. glutamicum*, the promoter *recA* (P*_recA_*) was fused to genes encoding for fluorescent proteins as optical readout. The derived DNA-stress biosensors like *P_recA__e2-crimson* enabled to monitor the SOS-response upon applying DNA-damage inducing agents (Nanda *et al*., 2014; Helfrich *et al*., 2015). As illustrated above, a broad portfolio of different biosensors targeting oxidative stress and DNA-stress are available. But, to the best of our knowledge, these biosensors have not been combined in one cell yet. Taking into account the complex linkage of different stress response systems such as oxidative DNA-damage, it becomes clear that novel biosensor concepts capable of targeting stresses in a multi-parametrical manner are required. Towards a first combinatorial biosensor concept, *C. glutamicum* was selected as a model system due to the availability of well-established biosensors such as the redox biosensor protein Mrx1-roGFP2 (Tung *et al*., 2019; Hartmann *et al*., 2020) and the transcriptional DNA-stress reporter system *P_recA__e2-crimson* (Nanda *et al*., 2014; Helfrich *et al*., 2015).

In this communication we show that the individual functionality of the biosensors Mrx1-roGFP2 and *P_recA__e2-crimson* were not affected when both biosensors were combined in *C. glutamicum*. By the use of the double biosensor system, we show that oxidative stress applied *via* artificial oxidants induces an oxidative stress response accompanied by a DNA-damage response. In absence of the antioxidant MSH, the underlying DNA-stress response was even more pronounced, indicating the linkage of oxidative stress and DNA-damage. We further show that DNA-damage alone initiated an oxidative redox shift, suggesting a direct link of DNA-damage and oxidative stress response in *C. glutamicum*. Furthermore, inhibition of cell wall synthesis in *C. glutamicum* resulted in an oxidative redox shift as well as DNA-damage response. This study demonstrates the compatibility of redox proteins based on roGFP2 with the fluorescent reporter E2-Crimson and therefore accelerates various biosensor combinations to facilitate physiological redox studies in the future.

## 2 Material and Methods

### 2.1 Strains, media and culture conditions

Bacterial strains and plasmids used in this study are listed in Table 1. *C. glutamicum* strains were pre-cultured in Lysogeny Broth (LB) medium (Bertani, 2004) at 30 °C as 5 mL or 50 mL cultures using cultivation tubes or shaker flasks, respectively. Prior to inoculating the main-culture, cells of an overnight culture were washed twice with 100 mM potassium phosphate buffer (PBS; 100 mM; pH 7.0). If not otherwise stated, *C. glutamicum* main cultures were grown in CGXII minimal medium (Graf *et al*., 2018). Growth was performed in 500 mL shaker flasks (50 mL cultures) or as 800 μL cultures using a BioLectorII system (m2p-labs, Baesweiler/DE) and 48-well Flowerplates (m2p-labs, Baesweiler/DE) at 1200 rpm and 30 °C as recently described (Hartmann *et al*., 2021). *E. coli* strains were pre-cultured in LB medium at 37 °C as 5 mL or 50 mL cultures. For protein production (Mrx1-roGFP2 biosensor protein or the fluorescent protein E2-Crimson), 50 mL LB media was inoculated to a final optical density of 0.1 at 600 nm (OD_600nm_). IPTG was added to a final concentration of 1 mM when the culture reached an OD_600nm_ between 0.6-0.8. Cells were cultured in presence of appropriate antibiotics using a final concentration of 50 μg/mL (kanamycin or ampicillin).

**Table 1:**
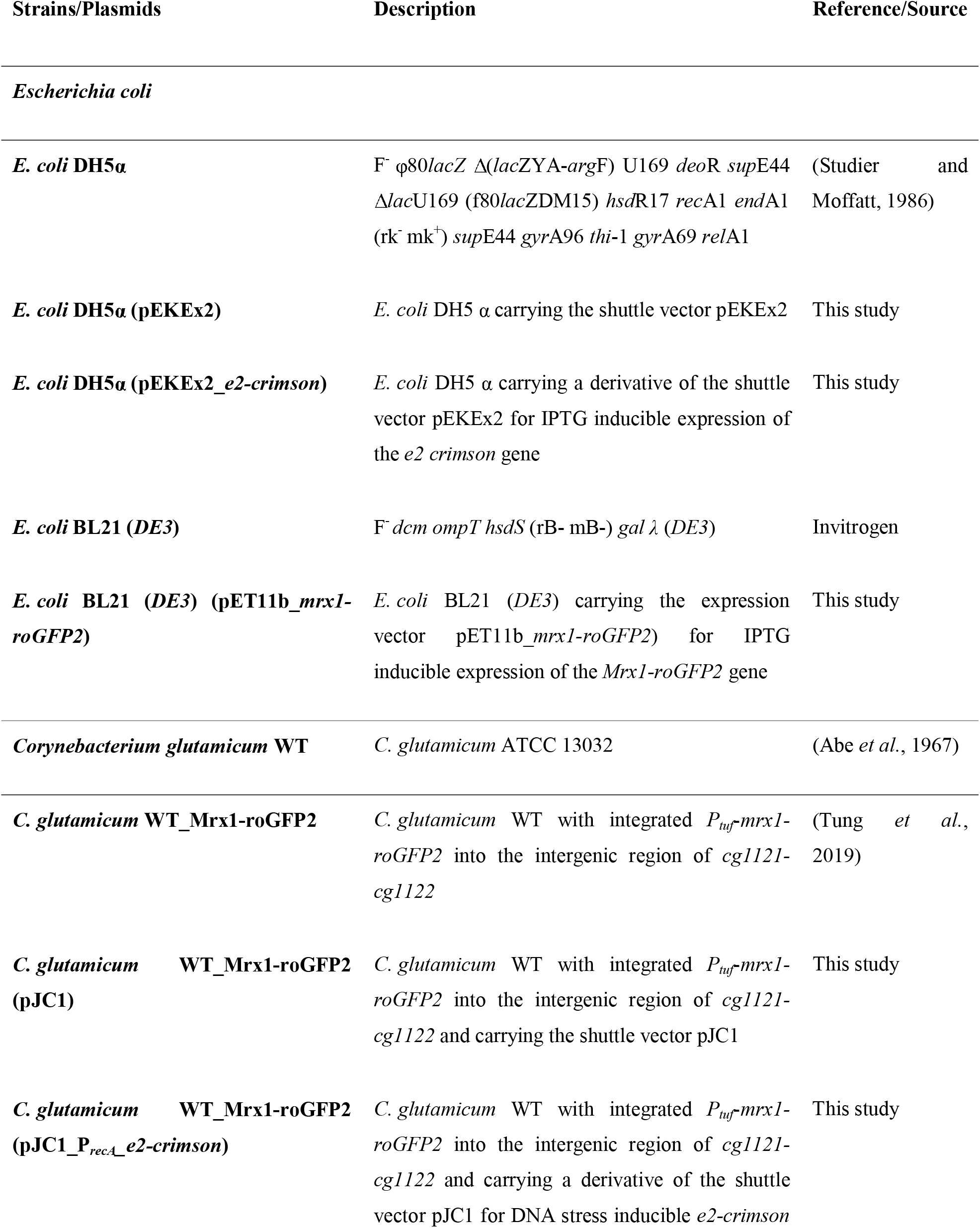
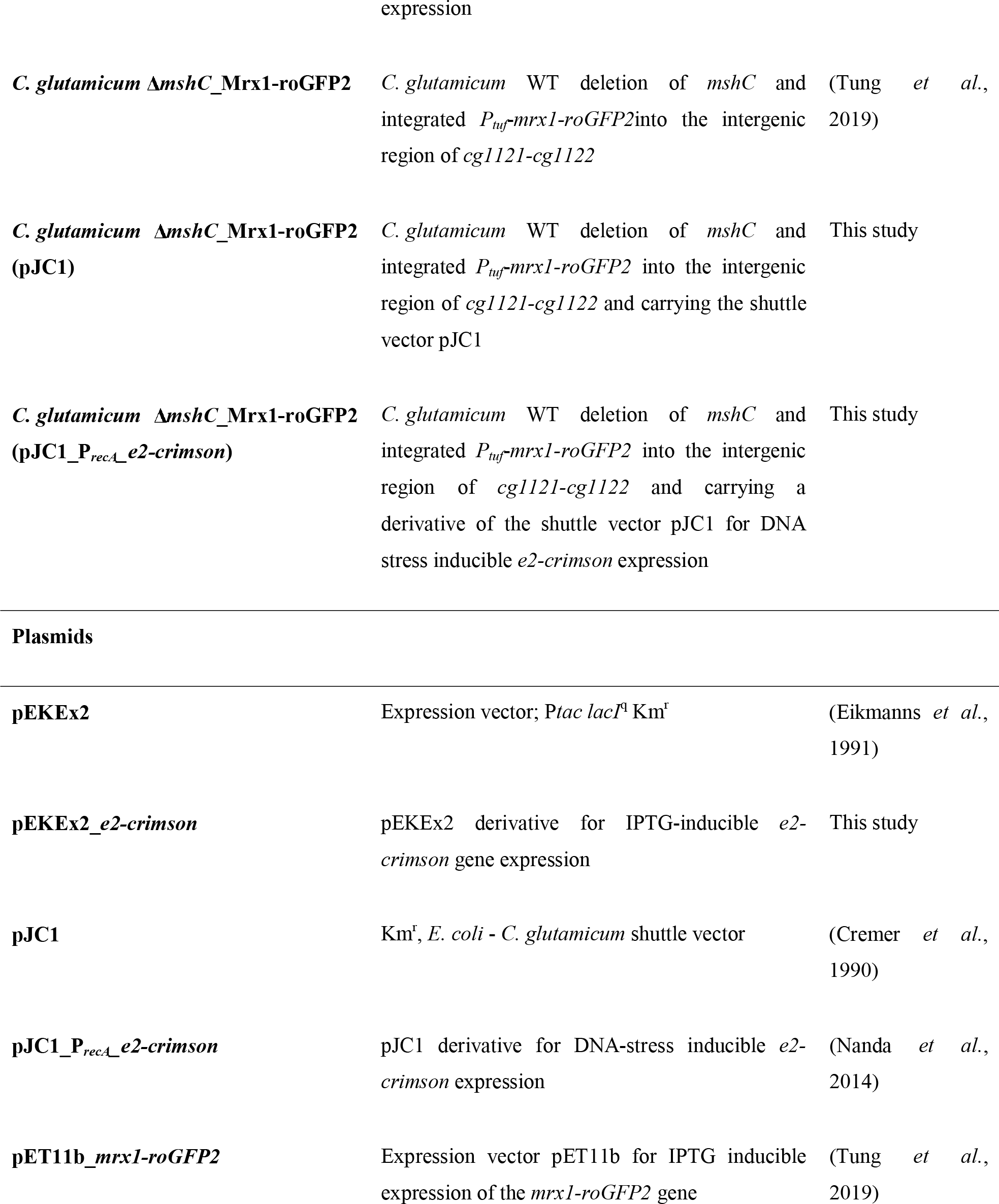
Bacterial strains and plasmids used in this study.

### 2.2 Construction of pEKEx2_*e2-crimson*

For the construction of the plasmid pEKEx2_*e2-crimson* (see Table 1), the gene *e2-crimson* encoding for the red fluorescent protein E2-Crimson was amplified from pJC1_P*_recA__e2-crimson* using the primer pair ggatccccgggtaccgagct<u>aaaggaggcccttcag</u>ATGGATAGCACTGAGAAC and acggccagtgaattcgagctCTACTGGAACAGGTGGTG (small letters: overhang, underlined letters: ribosomal binding site, capital letters: annealing part). The plasmid pEKE2 was linearized with *SacI* (New England Biolabs, U.S.A.) followed by assembling of the amplified gene *via* Gibson Assembly according to the manufacturer’s instructions (Gibson Assembly^®^ Master Mix, New England Biolabs, U.S.A.). Identity of the obtained plasmid pEKEx2_*e2-crimson* was verified *via* colony PCR.

### 2.3 *In-vitro* fluorescence analysis

Prior to *in-vitro* fluorescence analysis of the redox biosensor protein Mrx1-roGFP2 and the red fluorescent protein E2-Crimson, *E. coli* BL21 (*DE3*) (pET11b_*mrx1-roGFP2*) and *E. coli* DH5α (pEKEx2_*e2*-*crimson*) were pre-cultured in 5 mL LB medium, respectively. After six hours of cultivation, cells were transferred to 50 mL LB medium by adjusting an OD_600nm_ of 0.1. Cells were cultured until an OD_600nm_ of 0.6 - 0.8 was reached. Then, expression of the genes encoding for Mrx1-roGFP2 or E2-Crimson were induced by adding 1 mM (final concentration) IPTG to the respective cultures and cultivated further for 16 hours (37 °C and 150 rpm). Following that, cells were harvested and washed twice with PBS buffer (100 mM, pH 7.0). Finally, an OD_600nm_ of 40 was adjusted and crude cell extracts prepared as recently described (Hartmann *et al*., 2020; Hartmann *et al*., 2022). Cell debris were removed by centrifugation (12,000 rpm, 20 min.; 4 °C) and 180 μL of the supernatant transferred to black flat-bottomed 96-well microplates (Thermo Fisher Scientific, Germany). Fluorescence analysis was conducted in a plate reader device (SpectraMax iD3, Molecular Devices LLC, U.S.A). For fluorescence analysis using E2-Crimson, the supernatant (180 μL) obtained from the crude cell extracts of *E. coli* DH5 α (pEKEx2_*e2-crimson*) was supplemented with 20 μL of methyl-methanesulfonate (MMS), dithiothreitol (DTT), sodium hypochlorite (NaOCl), cumene hydroperoxide (CHP) using differently concentrated stock solution to adjust final concentrations of 350 mM; 100 mM (MMS), 20 mM; 10 mM (DTT), 70 mM; 5 mM (NaOCl), 100 mM; 50 mM (CHP) in the wells. As a control, PBS buffer only was added. After an incubation of 15 min at 37 °C and shaking, fluorescence intensity was recorded at 640 nm by setting an excitation wavelength of 600 nm. Fluorescence intensities of treated samples were compared to the PBS control as indicator if the fluorescence properties of E2-Crimson are negatively affected by respective chemicals. For analysis of the redox biosensor Mrx1-roGFP2, the supernatant (180 μL) obtained from the crude cell extracts of *E. coli* BL21 (*DE3*) (pET11b_*mrx1-roGFP2*) was supplemented with 20 μL of DTT (20 mM; final concentration) and NaOCl (70 mM; final concentration) to obtain fully reduced and oxidized controls, respectively. Then, 20 μL of MMS, PBS or supernatant obtained from crude cell extracts with E2-Crimson were added to verify the stability of the adjusted oxidation degree (OxD) of the biosensor protein Mrx1-roGFP2 *in-vitro* (Equation 1). Fluorescence measurements were conducted with a set emission intensity at 510 nm and excitation at 380 nm and 470 nm as recently described (Hartmann *et al*., 2020).

### 2.4 *In-vivo* fluorescence analysis

For *in-vivo* characterization of the DNA-stress reporter plasmid pJC1_P*_recA__e2-crimson*, *C. glutamicum* WT was transformed with the plasmid to obtain the reporter strain *C. glutamicum* WT (pJC1_P*_recA__e2-crimson*). As a control, pJC1 empty vector was introduced into *C. glutamicum* WT, resulting in the control strain *C. glutamicum* WT (pJC1). Functionality of the DNA-stress reporter plasmid was verified by performing micro-fermentations using a BioLectorII system as described above. After four hours of cultivation, the experiment was paused and different MMS concentrations (0 - 800 μg/mL) added to the respective wells. As a control, PBS buffer only was added. E2-Crimson signal was recorded using a RFP filter in the BioLectorII system. At the end of the experiment, samples were taken and prepared for subsequent *off-line* fluorescence analysis. Recorded E2-Crimson fluorescence intensities were normalized to an OD_600nm_ of 1 for both the DNA-stress reporter plasmid strain as well as empty vector control strain for all of the tested conditions.

For *in-vivo* characterization of *C. glutamicum* WT_Mrx1-roGFP2 (pJC1), *C. glutamicum* WT (pJC1_P*_recA__e2-crimson*), *C. glutamicum* Δ*mshC*_Mrx1-roGFP2 (pJC1) and *C. glutamicum ΔmshC* (pJC1_P*_recA__e2-crimson*), strains were cultivated in LB medium in presence of kanamycin at 30 °C and 150 rpm for 16 hours (50 mL medium; shaker flasks). To verify the functionality of the DNA-stress reporter plasmid, 100 μg/mL and 800 μg/mL MMS was added to the shaker flasks after four hours of cultivation and further cultivated for 16 hours. Following that, cells were harvested *via* centrifugation, washed twice with PBS buffer and an OD_600nm_ of 40 adjusted as described. Prior to fluorescence analysis, 200 μL of the cell suspension was transferred to black flat-bottomed 96-well microplates, and fluorescence intensity at 640 nm (Exc. 600 nm) recorded to verify the E2-Crimson intensity of the respective strains. Emission intensity was divided by 40 to obtain relative fluorescence units (RFLU) normalized to an OD_600nm_ of 1.

In order to determine the Mrx1-roGFP2 biosensor functionality, the calculation of Exc. 380 nm/470 nm (emission intensity 510 nm) was used to determine the biosensor ratio, which is known to increase upon oxidation and *vice versa* upon reduction of the biosensor probe as recently described in Hartmann *et al*., 2020 (Hartmann *et al*., 2020). For this, 180 μL of the cell suspension (OD_600nm_ of 40) was transferred to black-bottomed 96-well plates. Prior to fluorescence analysis, 20 μL of PBS as control and different final concentrations of DTT (0 - 20 mM) or NaOCl (0 - 70 mM) was adjusted for the cell suspensions in the wells in order to induce biosensor reduction or oxidation, respectively. The untreated sample was used to determine the OxD value (Equation 1) (Bhaskar *et al*., 2014) of the different samples and to further analyze the redox potential E_MSH_ in the different tested strain backgrounds using the Nernst equation as recently described (Bhaskar *et al*., 2014; Tung *et al*., 2019; Hartmann *et al*., 2020) (Equation 2).

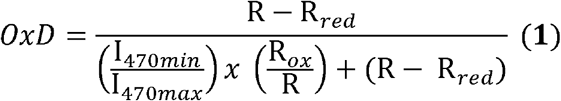

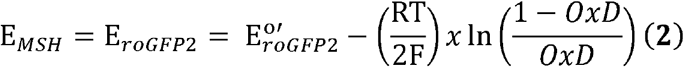

In equation 1, I_470min_ and I_470max_ represent the measured fluorescence intensities received for an excitation at 470 nm (Em. 510 nm) for fully oxidized and reduced biosensor probes, respectively. Moreover, R, R_red_, R_ox_ is the biosensor ratio Exc. 470 nm/380 nm determined for the sample (untreated), fully reduced or fully oxidized biosensor protein, respectively.

In order to determine the redox potential of MSH, the calculated OxD values, the standard midpoint redox potential of roGFP2 (*E*°’roGFP2= −280 mV) (Dooley *et al*., 2004), Faraday constant (F: 96,485 C mol^-1^) of electric charge per mole of electrons, the respective temperature in Kelvin (T: 303.15 K) and the universal gas constant (R: 8.314 J K^-1^ mol^-1^) were incorporated into Equation 2.

### 2.5 Growth experiment under artificially induced oxidative stress conditions using the oxidative stress and DNA-stress reporter strain

*C. glutamicum* WT (pJC1_P*_recA__e2-crimson*) and *C. glutamicum ΔmshC* (pJC1_P*_recA__e2-crimson*) were pre-cultured as mentioned before. Following that, 100 mL of minimal medium CGXII, supplemented with 1% (w/v) Glucose as sole carbon source, was inoculated in shaker flasks and growth monitored by measuring the OD_600nm_. After four hours of cultivation, the culture was divided into two separate cultures. Subsequently, 100 μL PBS buffer or NaOCl was added to obtain the untreated and the treated sample, respectively. In order to study the response (oxidative stress) as well as DNA-stress *via* the reporter plasmid pJC1_P*_recA__e2-crimson*, the production of the fluorescent protein E2-Crimson functions as optical output. Moreover, the cells need to maintain their viability to respond and adapt to the changed conditions and in turn to produce the fluorescent protein. To ensure that the viability is maintained, impedance flow cytometry measurements (IFC) were conducted as previously described (Hartmann *et al*., 2021). Based on this, 35 mM NaOCl was added in order to induce oxidative stress. Initially, after adding the artificial oxidant, the OxD value of the redox biosensor Mrx1-roGFP in the double sensor strain for both the treated as well as the untreated sample was measured as described in the section *in-vivo* fluorescence analysis. Moreover, E2-Crimson fluorescence signal was recorded for both samples as described. The fluorescence intensity was normalized to an OD_600nm_ of 1, and the emission intensity obtained for the treated sample divided by the intensity measured for the untreated sample in order to calculate the E2-Crimson ratio (E2-Crimson (treated/untreated)). Consequently, a value of >1 indicates a response of the transcriptional DNA-stress reporter plasmid, whereas a value of 1 is expected to be assessed in absence of any response by the DNA-stress reporter. The growth experiment was performed until the stationary phase was reached (after 10 hours).

### 2.6 Data analysis

Analysis of one-way variance (ANOVA) with Tukey’s test was used to assess differences of sensor signals derived from *in-vitro* and *in-vivo* analysis. Differences were considered statistically significant when *p* < 0.05.

## 3 Results and Discussion

### 3.1 E2-Crimson is a suitable fluorescent reporter protein in oxidized environments

Prior to combining two genetically encoded biosensors in one cell, several aspects need to be considered such as:

i. the excitation and emission wavelengths of the applied fluorescent proteins should not interfere with each other
ii. the properties of fluorescent proteins should not be affected by chemicals required to apply stresses required for physiological studies and characterization purposes

The redox biosensor protein Mrx1-roGFP2 was intensively characterized *in-vitro* and *in-vivo* (Bhaskar *et al*., 2014; Tung *et al*., 2019; Hartmann *et al*., 2020) and recently successfully applied in *C. glutamicum* for assessing the redox environment in different strain backgrounds (Tung *et al*., 2019; Hartmann *et al*., 2020). Thus, this biosensor protein was considered as an appropriate fluorescent reporter protein towards assessing oxidative stress levels in *C. glutamicum*. The fluorescent protein roGFP2 consists of Cys residues that form a disulfide bond upon oxidation, resulting in a ratiometric change of two excitation maxima (Exc. 380 nm/ 470 nm) (Hartmann *et al*., 2020) with a single emission wavelength at 510 nm (green spectra) (Schwarzländer *et al*., 2016). Accordingly, the fluorescence properties of Mrx1-roGFP2 should allow its combination with a second fluorescent protein in the red spectra. Therefore, a recently developed reporter system based on the RecA-dependent transcription of a gene encoding for the red fluorescent protein E2-Crimson (Exc. 600 nm/ Em. 640 nm) was selected as promising DNA-stress reporter towards monitoring DNA-damage in *C. glutamicum* (Nanda *et al*., 2014).

To test the stability of the E2-Crimson fluorescence in presence of oxidants/reductants, cell-free extracts derived from *E. coli* DH5 *α* (pEKEx2_*e2-crimson*) were incubated for 15 min with various amounts of DTT (reducing agent), NaOCl (oxidizing agent), and CHP (oxidizing agent) prior to performing the fluorescence analysis (Fig. 1a). The fluorescence measurements revealed that the emission intensity at 640 nm (Exc. 600 nm) for the control sample (+ PBS buffer) was between 2.7 x 10^5^ – 2.8 x 10^5^ FLU, which is depicted as dashed horizontal line in Fig. 1a. The addition of DTT, NaOCl or CHP did not lead to a significant change of the measured fluorescence intensity when compared to the PBS control (Fig. 1a). This indicates the stability of E2-Crimson and by this its capability as appropriate optical readout for transcriptional reporter constructs in an oxidized environment. Moreover, E2-Crimson maintains its fluorescence properties when exposed to MMS, a common alkylating agent used to induce DNA-damage. This confirms that the DNA-damage inducing agent MMS does not affect fluorescence of the reporter protein E2-Crimson, which shows this combination is well-suited to investigate DNA-stress response.

**Figure 1:**
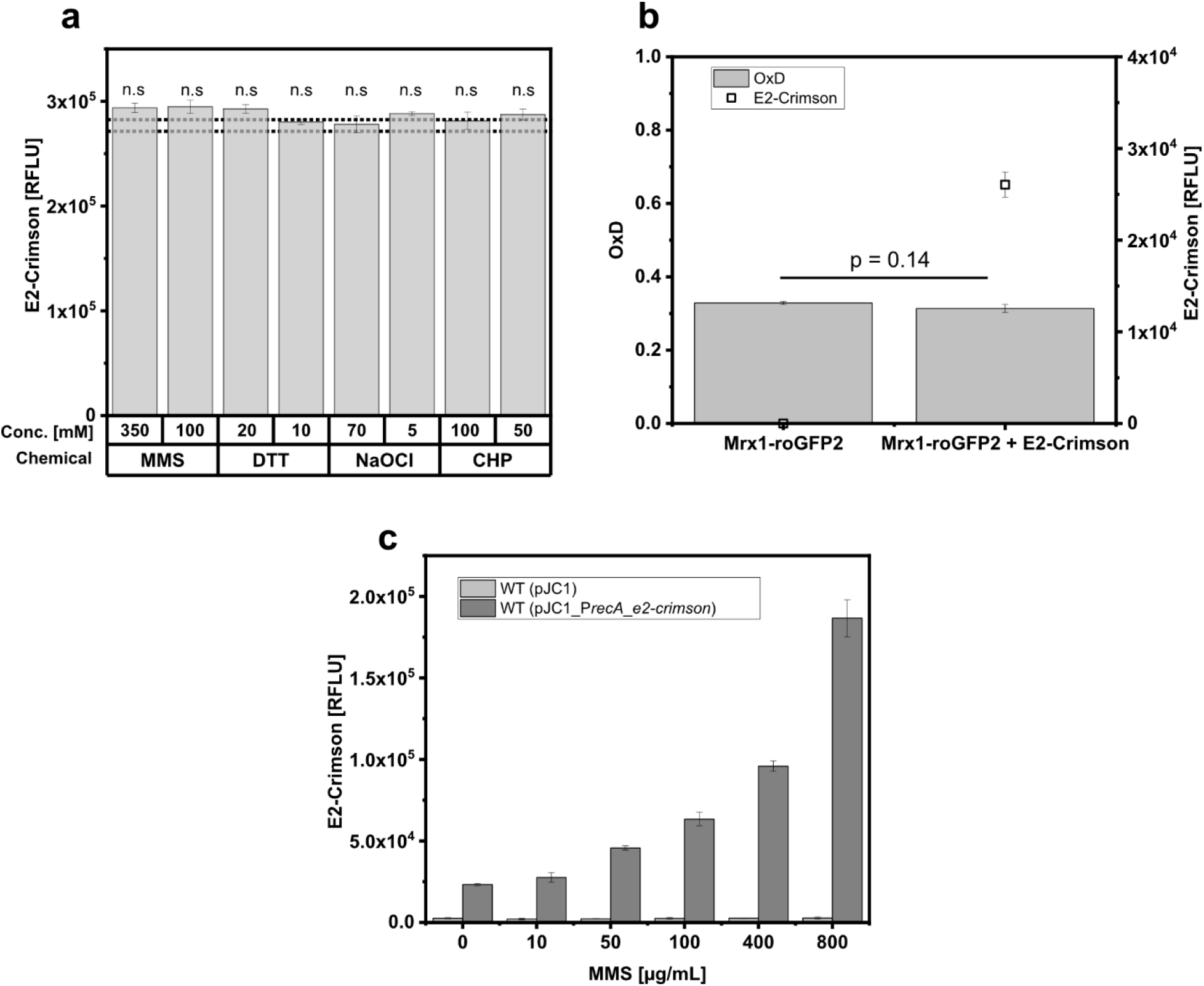
E2-Crimson fluorescence signal stability upon addition of methyl-methanesulfonate (MMS), dithiothreitol (DTT), sodium-hypochlorite (NaOCl) and cumene-hydroperoxide (CHP) using cell extracts of *E. coli* (pEKEx2_*e2-crimson*) **(A)** and stability of the oxidation degree (OxD) of Mrx1-roGFP2 in absence and presence of the fluorescent protein E2-Crimson by using cell extracts of *E. coli* (pET11b_*mrx1-roGFP2*) mixed with *E. coli* DH5 α (pEKEx2) or with *E. coli* DH5 α (pEKEx2_*e2-crimson*) cell extracts, respectively **(B)**. Relative E2-Crimson fluorescence intensitiesafter 22 hours of micro-fermentation conducted with different MMS concentrations for *C. glutamicum* WT harboring the empty vector control (pJC1) and the transcriptional DNA-damage biosensor pJC1_P*_recA__e2-crimson* **(C)**. Mrx1-roGFP2 fluorescence signal was measured by recording the emission intensity at 510 nm upon excitation at 380 nm and 470 nm. The ratiometric signal was calculated by dividing the former emission intensity by the latter followed by calculating the OxD values according to Equation 1 (Material and Methods). E2-Crimson fluorescence intensity was recorded at 640 nm upon an excitation at 600 nm. All fluorescence measurements were conducted in black flat-bottomed 96-well microtiter plates using a plate reader device (SpectraMaxiD3). Error bars represent standard deviation from at least three replicates. Statistical analysis was performed via One-Way-ANOVA followed by a Tukey’s test (^ns^ p > 0.05).

To further test the compatibility of Mrx1-roGFP2 and E2-Crimson *in-vitro*, a cell free extract containing Mrx1-roGFP2 was generated from an *E. coli* BL21 (*DE3*) (pET11b_*mrx1-roGFP2*) culture. The oxidation degree (OxD) of this extract was adjusted by titration with DTT or NaOCl to approximately 0.3 and then the OxD-adjusted extract split into several batches of 180 μL. Batches were mixed with either 20 μL cell-free extract containing E2-Crimson (prepared from an *E. coli* DH5 α (pEKEx2_*e2-crimson*) culture) or 20 μL of cell-free extract without additional fluorescent protein (prepared from an *E. coli* DH5 α (pEKEx2) culture). Subsequently, the fluorescence for Mrx1-roGFP2 and E2-Crimson was analyzed. As depicted in Fig. 1b, the addition of E2-Crimson resulted in an increase of fluorescence units (2.6 x 10^4^ ± 1373 FLU, excitation at 600 nm, emission at 640 nm) when compared to the control sample. In contrast, the OxD values determined for the redox biosensor Mrx1-roGFP2 were measured to be 0.33 ± 0.00 and 0.31 ± 0.01 for samples in absence and presence of E2-Crimson, respectively (Fig. 1b). Thus, it can be concluded that the presence of E2-Crimson does not cause a significant OxD difference (p = 0.14) and thus not affecting the functionality of Mrx1-roGFP2. Taken together the *in-vitro* results show that E2-Crimson is an appropriate fluorescent protein to be combined with the redox biosensor protein Mrx1-roGFP2 due to its stability in an oxidized environment and the fact that it does not affect the oxidation state of sensor protein.

### 3.2 Characterization of the response of the DNA-damage sensor P*_recA__e2-crimson*

In order to characterize the response of the reporter system P*_recA__e2-crimson* to the DNA-damaging agent MMS, the *C. glutamicum* WT was equipped with the plasmid pJC1_P*_recA__e2-crimson* or the empty vector pJC1 resulting in the reporter strain *C. glutamicum* WT (pJC1_P*_recA__e2-crimson*) and the control strain *C. glutamicum* WT (pJC1), respectively. To analyze the biosensor signal during cultivation at various MMS concentrations, the strains were cultured using a BioLectorII system (Hartmann *et al*., 2021). After four hours of cultivation, the cultivation was paused and 5 μL MMS of differently concentrated stock solutions were added to adjust different final concentrations in the cultivation wells. The E2-Crimson signal of the DNA-damage reporter P*_recA__e2-crimson* increased in a MMS concentration dependent manner (Fig. S1), which indicates the functionality of the DNA-stress reporter plasmid pJC1_P*_recA__e2-crimson*. Endpoint measurements conducted in a plate reader device confirmed the dose dependent impact of MMS on measured E2-Crimson levels in the reporter strains (Fig. 1c). In opposite, the empty vector control did not show any response with respect to the measured endpoint fluorescence signals (Fig. 1c). Taken together the results confirmed that the P*_recA__e2-crimson* is a functional DNA-stress reporter for use in *C. glutamicum*.

### 3.3 Combinatorial use of Mrx1-roGFP2 and P*_recA__e2-crimson* allows to simultaneously measure oxidative stress and DNA-stress levels in *C. glutamicum*

To test the combination of the protein based redox biosensor Mrx1-roGFP2 with the transcriptional DNA-stress sensor P*_recA__e2-crimson in-vivo, C. glutamicum* WT_Mrx1-roGFP2 and the MSH-deficient mutant strain *C. glutamicum* Δ*mshC*_Mrx1-roGFP2, which both harbor a genome integrated gene for Mrx1-roGFP2 (Tung *et al*., 2019), were transformed with the plasmid pJC1_P*_recA__e2-crimson*, resulting in the strains *C. glutamicum* WT_Mrx1-roGFP2 (pJC1_P*_recA__e2-crimson*) and *C. glutamicum* Δ*mshC*_Mrx1-roGFP2 (pJC1_P*_recA__e2-crimson*). In addition, the empty-plasmid control strains *C. glutamicum* WT_Mrx1-roGFP2 (pJC1) and *C. glutamicum* Δ*mshC*_Mrx1-roGFP2 (pJC1) were generated. To test the P*_recA__e2-crimson* biosensor-response, the strains were cultivated in LB-medium in presence of 100 μg/mL MMS, 800 μg/mL MMS and absence of MMS for 16 hours. Prior to fluorescence analysis, cells were harvested and washed twice with PBS buffer and re-suspended in the same buffer to a final OD_600nm_ of 1 for all samples. As expected, fluorescence measurements revealed a response of the DNA-stress reporter P*_recA__e2-crimson*, indicated by increased emission intensities measured at 640 nm (Exc. 600 nm) for the double sensor strains *C. glutamicum* WT_Mrx1-roGFP2 (pJC1_P*_recA__e2-crimson*) and *C. glutamicum* Δ*mshC*_Mrx1-roGFP2 (pJC1_P*_recA__e2-crimson*) (Fig. 2a, b). In detail, for *C. glutamicum* WT_Mrx1-roGFP2 (pJC1_P*_recA__e2-crimson*) the relative fluorescence units (RFLU) were measured to be 1.85 x 10^4^ ± 458, 4.17 x 10^4^ ± 2063 and 1.28 x 10^5^ ± 5442 for the approach without any supplements, 100 μg/mL MMS and 800 μg/mL MMS, respectively (Fig. 2a). In contrast, E2-Crimson signals for the empty vector control strain *C. glutamicum* WT_Mrx1-roGFP2 (pJC1) of 374 ± 4, 1685 ± 5 and 1776 ± 5 were detected for the approach without any supplements, 100 μg/mL MMS, and 800 μg/mL MMS, respectively, which are lower values when compared to the strains carrying the reporter plasmid (Fig. 2a). For *C. glutamicum* Δ*mshC*_Mrx1-roGFP2 (pJC1_P*_recA__e2-crimson*) values of 1.80 x 10^4^ ± 934 RFLU, 5.08 x 10^4^ ± 4155 RFLU, and 1.32 x 10^5^ ± 5529 RFLU were measured for the approaches without MMS, 100 μg/mL and 800 μg/mL MMS, respectively (Fig. 2b). These values are in the same range as measured for *C. glutamicum* WT_Mrx1-roGFP2 (pJC1_P*_recA__e2-crimson*). Similar to the *C. glutamicum* WT_Mrx1-roGFP2 (pJC1), the empty vector control strain *C. glutamicum* Δ*mshC*_Mrx1-roGFP2 (pJC1) revealed fluorescence intensities of 763 ± 5 RFLU, 1812 ± 6 RFLU and 2756 ± 24 RFLU for the approach without MMS, 100 μg/mL and 800 μg/mL MMS, respectively (Fig. 2b).

**Figure 2:**
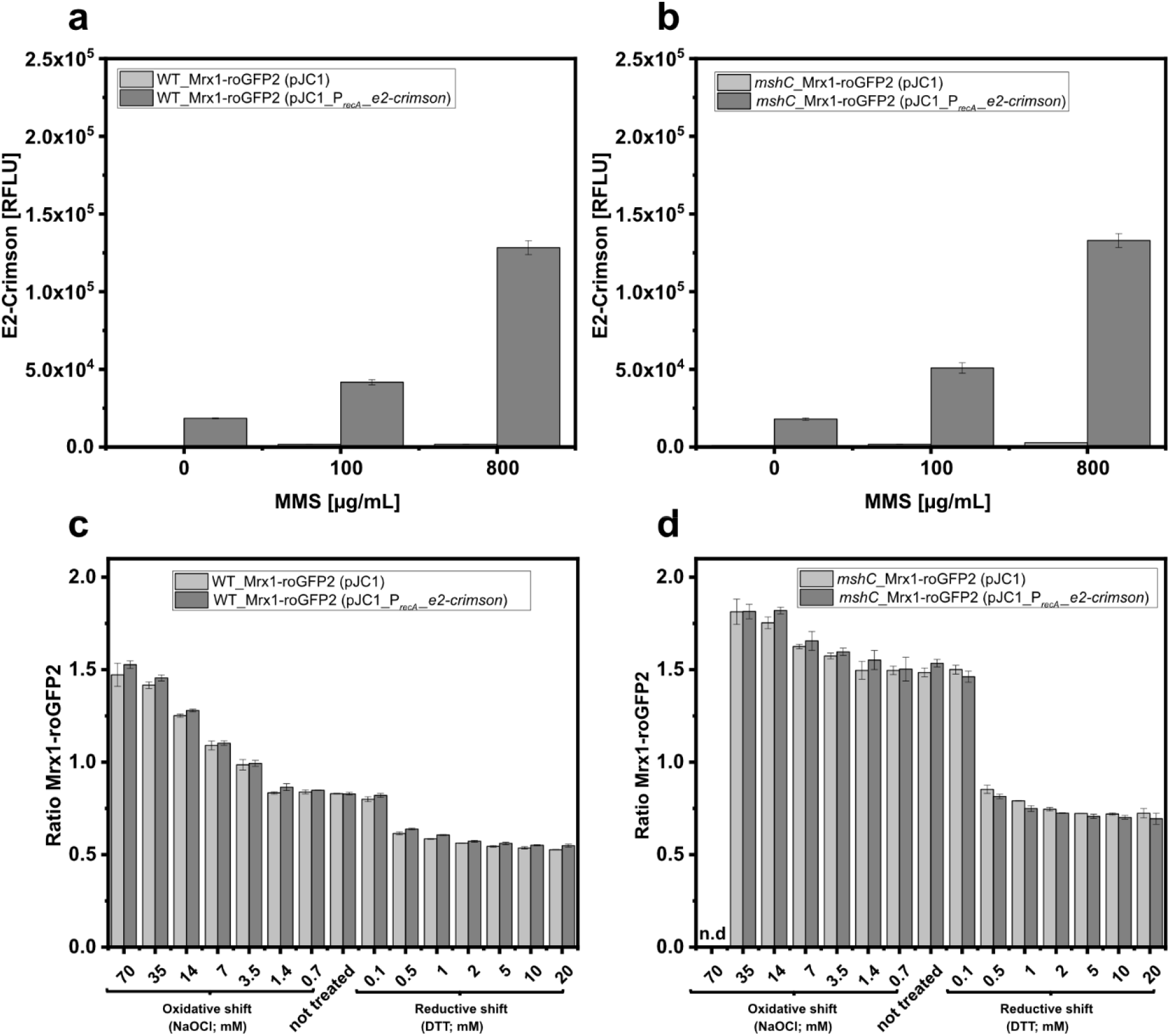
Relative E2-Crimson fluorescence intensities as indicator of the DNA-damage response after 24 hours cultivation conducted in shaker flasks and in presence of different methyl-methanesulfonate (MMS) concentrations using the double biosensor strains *C. glutamicum* WT_Mrx1-roGFP2 (pJC1_*P_recA__e2-crimson*) and the empty vector control *C. glutamicum* WT_Mrx1-roGFP2 (pJC1) **(A)** *C. glutamicum* Δ*mshC*_Mrx1-roGFP2 (pJC1_*P_recA__e2-crimson*) and the empty vector control *C. glutamicum* Δ*mshC*_Mrx1-roGFP2 (pJC1) **(B)**. Oxidative (sodium-hypochlorite (NaOCl)) and reductive shift (dithiothreitol (DTT)) of the redox biosensor protein Mrx1-roGFP2 measured using cultures of the double biosensor strain WT_Mrx1-roGFP2 (pJC1_*P_recA__e2-crimson*) and the empty vector strain WT_Mrx1-roGFP2 (pJC1) **(C)** as well as the mycothiol deficient mutant Δ*mshC*_Mrx1-roGFP2 (pJC1_*P_recA_e2-crimson*) and the empty vector control Δ*mshC*_Mrx1-roGFP2 (pJC1) **(D)**. Mrx1-roGFP2 fluorescence signal was measured by recording the emission intensity at 510 nm upon excitation at 380 nm and 470 nm. The ratiometric signal was calculated by dividing the former emission intensity by the latter (Material and Methods). E2-Crimson fluorescence intensity was recorded at 640 nm upon an excitation at 600 nm. All fluorescence measurements were conducted in black flat-bottomed 96-well microtiter plates using a plate reader device (SpectraMaxiD3). Error bars represent standard deviation from at least three replicates.

Besides verifying the functionality of the DNA-stress reporter P*_recA__e2-crimson*, the functionality of the redox biosensor protein Mrx1-roGFP2 was tested in the double biosensor strains. For this purpose, cell suspensions of *C. glutamicum* WT_Mrx1-roGFP2 (pJC1_P*_recA__e2-crimson*), *C. glutamicum* Δ*mshC*_Mrx1-roGFP2 (pJC1_P*_recA__e2-crimson*), *C. glutamicum* WT_Mrx1-roGFP2 (pJC1), and *C. glutamicum* Δ*mshC*_Mrx1-roGFP2 (pJC1) were cultivated in absence of MMS. The fluorescence properties of Mrx1-roGFP2 in these suspensions were analyzed after causing a reductive or oxidative shift by addition of differently concentrated DTT or NaOCl stock solutions. For all four strains, the addition of NaOCl resulted in an increase of the fluorescence ratio detected for the Mrx1-roGFP2 biosensor (Fig. 2c, d), whereas DTT triggered the *vice versa* response of the biosensor fluorescence ratios (Fig. 2c, d). In detail, the initial biosensor ratio (untreated) was measured to be 0.83 ± 0.00 and 0.83 ± 0.01 for *C. glutamicum* WT_Mrx1-roGFP2 (pJC1) and *C. glutamicum* WT_Mrx1-roGFP2 (pJC1_P*_recA__e2-crimson*), respectively (Fig. 2c). Upon inducing an oxidative shift, the biosensor ratio increased and was measured to be 1.47 ± 0.06 and 1.53 ± 0.02 for the *C. glutamicum* WT_Mrx1-roGFP2 (pJC1) and *C. glutamicum* WT_Mrx1-roGFP2 (pJC1_P*_recA__e2-crimson*), respectively (Fig. 2c). In contrast, applying DTT as reducing agent resulted in a final biosensor ratio of 0.53 ± 0.01 and 0.55 ± 0.01 for the *C. glutamicum* WT_Mrx1-roGFP2 (pJC1) and *C. glutamicum* WT_Mrx1-roGFP2 (pJC1_P*_recA__e2-crimson*), respectively (Fig. 2c).

In opposite, the initial (untreated) biosensor ratio for the MSH-deficient mutant strain was higher with measured biosensor ratios of 1.49 ± 0.02 and 1.53 ± 0.02 for the *C. glutamicum* Δ*mshC*_Mrx1-roGFP2 (pJC1) and *C. glutamicum* Δ*mshC*_Mrx1-roGFP2 (pJC1_P*_recA__e2-crimson*) strains, respectively (Fig. 2d). Upon the addition of NaOCl, the Mrx1-roGFP2 biosensor ratio just slightly increased to a maximum ratio of 1.81 ± 0.07 (*C. glutamicum* Δ*mshC*_Mrx1-roGFP2 (pJC1)) and 1.82 ± 0.04 (*C. glutamicum* Δ*mshC*_Mrx1-roGFP2 (pJC1_P*_recA__e2-crimson*)) (Fig. 2d). The biosensor was fully oxidized upon applying 35 mM NaOCl and a further increase to 70 mM might result in an over-oxidation, which cannot be considered as reliable and thus was not assessed for the MSH-deficient mutant strain. The addition of DTT as reducing agent resulted in a fast decrease of the biosensor ratio and finally was determined to be 0.72 ± 0.02 and 0.69 ± 0.03 for *C. glutamicum* Δ*mshC*_Mrx1-roGFP2 (pJC1) and *C. glutamicum* Δ*mshC*_Mrx1-roGFP2 (pJC1_P*_recA__e2-crimson*), respectively (Fig. 2d). Slightly shifted biosensor ratios towards higher values were observed for both a fully reduced as well as fully oxidized biosensor protein in the MSH-deficient mutant background when compared to the wild type strains (for both the empty vector as well as DNA-stress reporter plasmid carrying strains). This shift, however, was also reported previously for the Mrx1-roGFP2 biosensor strains not carrying any plasmid and thus is not caused by introducing the plasmids to these redox biosensor strains (Hartmann *et al*., 2020).

In a next step the OxD value of the biosensor protein Mrx1-roGFP2 and MSH redox potential was calculated from the Mrx1-roGFP2 biosensor values, as previously described (Bhaskar *et al*., 2014; Tung *et al*., 2019; Hartmann *et al*., 2020). As shown in Table 2, the initial redox environment (untreated) was measured to be −284 ± 1 (OxD = 0.42 ± 0.01) and 286 ± 1 (OxD = 0.38 ± 0.01) for the *C. glutamicum* WT_Mrx1-roGFP2 (pJC1) and *C. glutamicum* WT_Mrx1-roGFP2 (pJC1_P*_recA__e2-crimson*), respectively. In contrast, the redox environment of the MSH-deficient mutant strains was measured to be more oxidized with redox potentials of −262 ± 2 (OxD = 0.80 ± 0.02) and −260 ± 2 (OxD = 0.82 ± 0.01) for the *C. glutamicum* Δ*mshC*_Mrx1-roGFP2 (pJC1) and *C. glutamicum* Δ*mshC*_Mrx1-roGFP2 (pJC1_P*_recA__e2-crimson*), respectively. For the strains *C. glutamicum* WT_Mrx1-roGFP2 and *C. glutamicum* Δ*mshC*_Mrx1-roGFP2), redox potentials of −280 ± 2 (OxD = 0.49 ± 0.04) and −255 ± 7 (OxD = 0.86 ± 0.05) were determined in absence of any externally applied stress in a previous study (Hartmann *et al*., 2020) (Table 2). This indicates, that the presence of the empty vector pJC1 as well as the reporter plasmid pJC1_P*_recA__e2-crimson* do not affect the biosensor readout as well as the cellular physiology with respect to the redox environment.

**Table 2:**
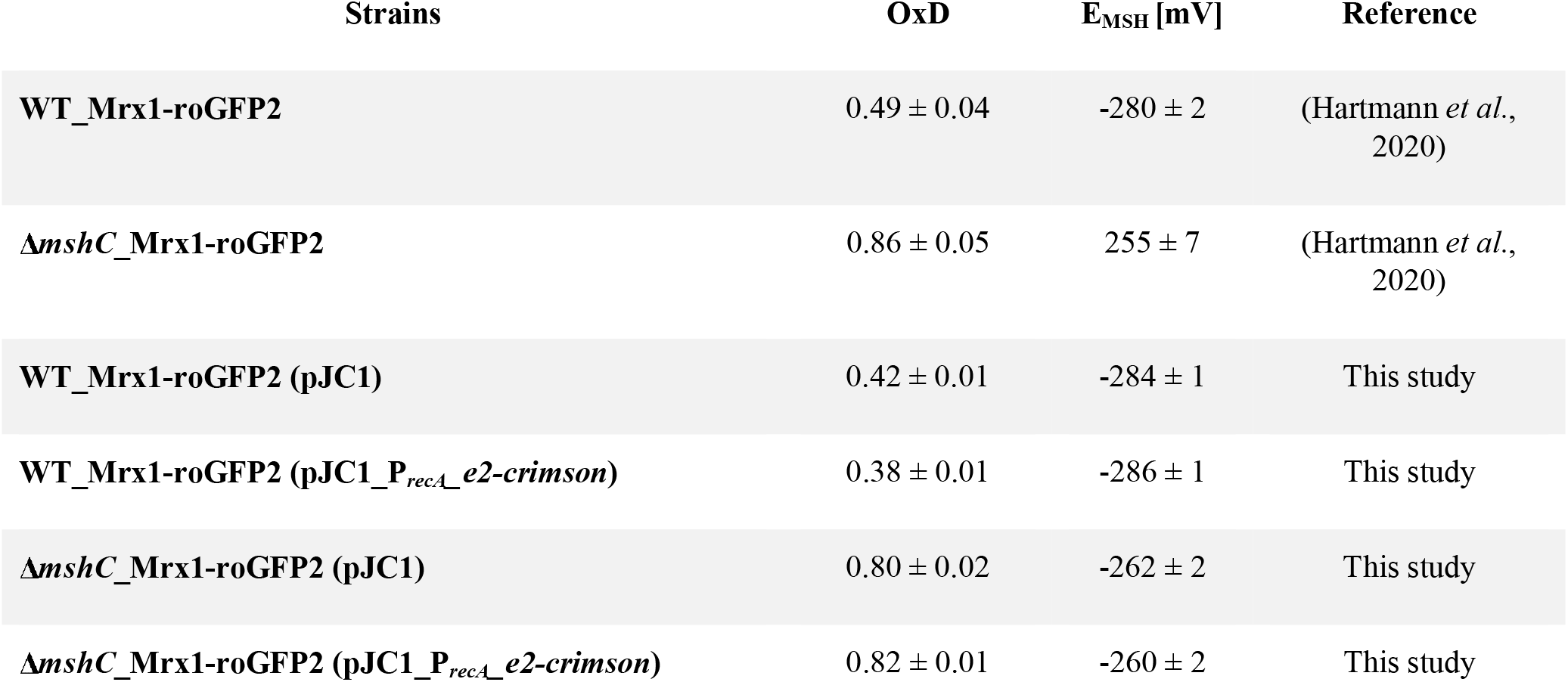
Oxidation degree (OxD) and corresponding redox potential for mycothiol (E_MSH_) determined in *C. glutamicum* WT and *C. glutamicum ΔmshC* harboring the genome integrated redox biosensor Mrx1-roGFP2 and additionally either the empty vector pJC1 or the DNA- stress reporter plasmid pJC1_P*_recA__e2-crimson*. Redox potential was determined *via* the Nernst equation. Data correspond to average values and respective standard deviation from at least three replicates (n=3).

| Strains | OxD | $E_{MSH}$ [mV] | Reference |
| --- | --- | --- | --- |
| WT_Mrx1-roGFP2 | $0.49 \pm 0.04$ | $-280 \pm 2$ | (Hartmann *et al.*, 2020) |
| $\Delta mshC$_Mrx1-roGFP2 | $0.86 \pm 0.05$ | $255 \pm 7$ | (Hartmann *et al.*, 2020) |
| WT_Mrx1-roGFP2 (pJC1) | $0.42 \pm 0.01$ | $-284 \pm 1$ | This study |
| WT_Mrx1-roGFP2 (pJC1_P<sub>recA</sub>_e2-crimson) | $0.38 \pm 0.01$ | $-286 \pm 1$ | This study |
| $\Delta mshC$_Mrx1-roGFP2 (pJC1) | $0.80 \pm 0.02$ | $-262 \pm 2$ | This study |
| $\Delta mshC$_Mrx1-roGFP2 (pJC1_P<sub>recA</sub>_e2-crimson) | $0.82 \pm 0.01$ | $-260 \pm 2$ | This study |

Taken together, the results show that the DNA-stress reporter plasmid pJC1_P*_recA__e2-crimson* is functional in the redox biosensor strains *C. glutamicum* WT_Mrx1-roGFP2 as well as *C. glutamicum* Δ*mshC*_Mrx1-roGFP2. Furthermore, the Mrx1-roGFP2 biosensor functionality is not affected in presence of the reporter plasmid when compared to empty vector controls and redox biosensor strains not harboring any plasmid (Hartmann *et al*., 2020). As a consequence, the two strains *C. glutamicum* WT_Mrx1-roGFP2 (pJC1_P*_recA__e2-crimson*) and *C. glutamicum* Δ*mshC*_Mrx1-roGFP2 (pJC1_P*_recA__e2-crimson*) are well suited to study the impact of oxidative stress on DNA-stress induction in *C. glutamicum*.

### 3.4 Oxidative stress induces the DNA-stress response in *C. glutamicum*

To study the impact of induced oxidative stress (measured *via* the redox biosensor Mrx1-roGFP2) on DNA-damage (measured *via* the P*_recA__e2-crimson* biosensor) the two strains *C. glutamicum* WT_Mrx1-roGFP2 (pJC1_P*_recA__e2-crimson*) and *C. glutamicum* Δ*mshC*_Mrx1-roGFP2 (pJC1_P*_recA__e2-crimson*) were cultivated in 100 mL of CGXII minimal medium with 1% (w/v) glucose as a sole carbon source for four hours prior to splitting the cultures in two 50 mL cultures. Afterwards, PBS buffer was added to one of the two obtained cultures of each strain (referred to as “untreated”), whereas NaOCl was added to the other culture (referred to as “treated”).

Prior to studying the interplay of oxidative stress and DNA-damage, the cultures viability was measured in regular intervals after applying oxidative stress using impedance flow cytometry (IFC). This allows one to ensure that the cultures can adapt to the harsh conditions rather than losing their cell viability and integrity and by this the capability of producing the reporter protein (E2-Crimson), as recently demonstrated for a non-growing but glutamate producing *C. glutamicum* culture (Hartmann *et al*., 2021).

Relative viabilities were calculated by dividing the viability of the treated culture by the untreated culture. Hereby, 1 refers to a value for which the determined viability of the treated sample was equal or higher when compared to the treated culture. The relative viability of *C. glutamicum* WT_Mrx1-roGFP2 (pJC1_P*_recA__e2-crimson*) and *C. glutamicum* Δ*mshC*_Mrx1-roGFP2 (pJC1_P*_recA__e2-crimson*) cultures were not affected three hours after NaOCl has been added (Fig. 3a, b). Following that, the relative viability slightly decreased in both cultures to around 80% and 60% for *C. glutamicum* WT_Mrx1-roGFP2 (pJC1_P*_recA__e2-crimson*) (Fig. 3a) and *C. glutamicum* Δ*mshC*_Mrx1-roGFP2 (pJC1_P*_recA__e2-crimson*) (Fig. 3b) cultures, respectively. A higher loss of cell viability for *C. glutamicum* Δ*mshC*_Mrx1-roGFP2 (pJC1_P*_recA__e2-crimson*) indicates their susceptibility towards oxidative stress and probably irreversible oxidative damages. But, still, the majority of cells in the culture remained viable (>50%) until entering the stationary growth phase (Fig. 3a, b). Consequently, the applied doses of NaOCl were considered as adequate towards studying the stress-response in *C. glutamicum*.

**Figure 3:**
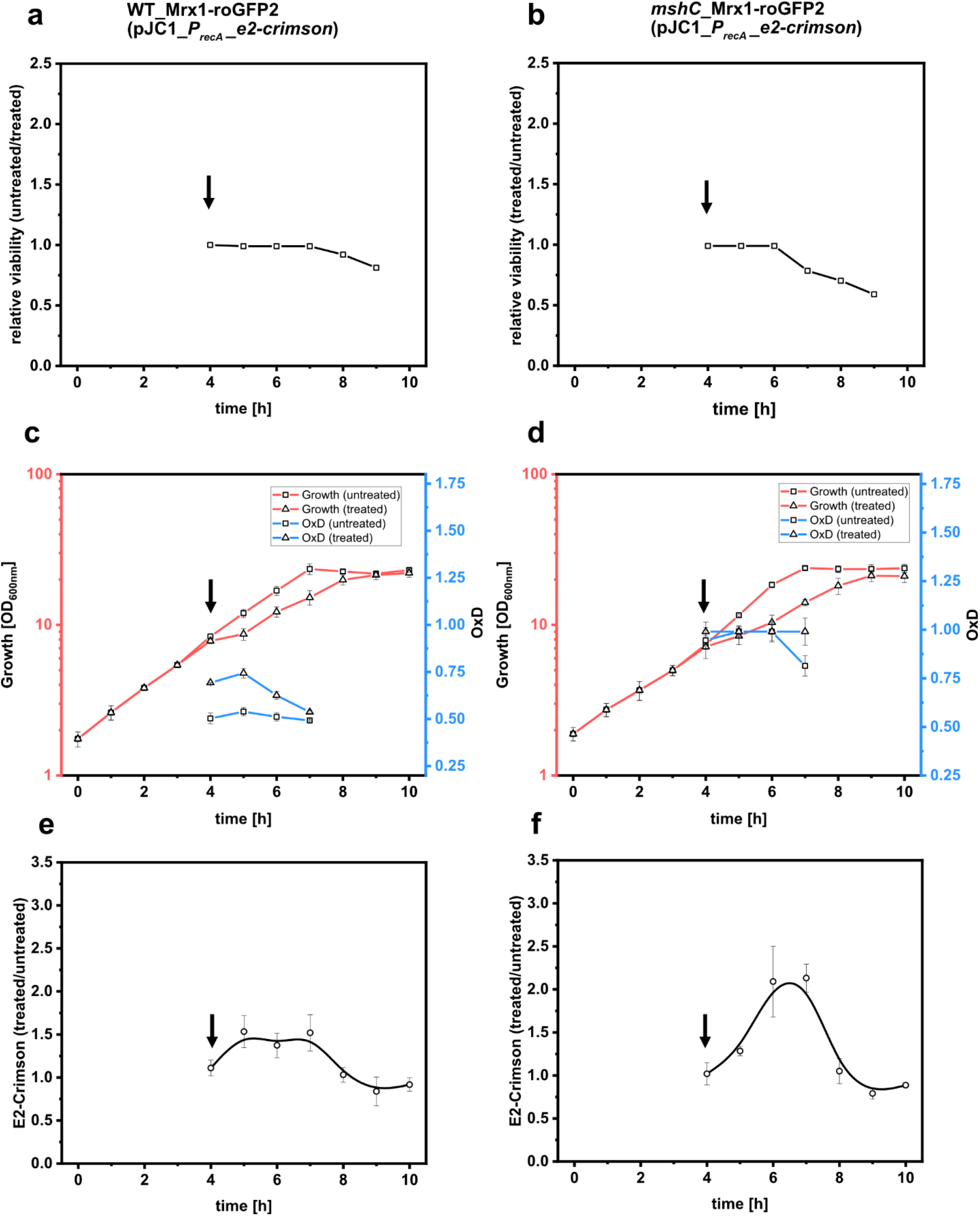
Relative viability determined by dividing the viable cells for the treated samples by the untreated samples for *C. glutamicum* WT_Mrx1-roGFP2 (pJC1_*P_recA__e2-crimson*) **(A)** and *C. glutamicum* Δ*mshC*_Mrx1-roGFP2 (pJC1_*P_recA__e2-crimson*) **(B)**. Monitoring of growth and oxidative stress in *C. glutamicum* WT_Mrx1-roGFP2 (pJC1_*P_recA__e2-crimson*) **(C)** and *C. glutamicum* Δ*mshC*_Mrx1-roGFP2 (pJC1_*P_recA__e2-crimson*) **(D)** *via* the redox sensor protein Mrx1-roGFP2 during shaker flask cultivations for both treated (NaOCl) and untreated (buffer control) samples. Ratio of the relative E2-Crimson fluorescence intensity determined by dividing the fluorescence intensity measured for the treated samples by the untreated samples of WT_Mrx1-roGFP2 (pJC1_*P_recA__e2-crimson)* **(E)** and Δ*mshC*_Mrx1-roGFP2 (pJC1_*P_recA__e2-crimson*) **(F)**. Mrx1-roGFP2 fluorescence signal was measured by recording the emission intensity at 510 nm upon excitation at 380 nm and 470 nm. The ratiometric signal was calculated by dividing the former emission intensity by the latter followed by calculating the OxD value according to equation1 (Material and Methods). E2-Crimson fluorescence intensity was recorded at 640 nm upon an excitation at 600 nm. All fluorescence measurements were conducted in black flat-bottomed 96-well microtiter plates using a plate reader device (SpectraMaxiD3). Viability (viable cells*mL^-1^*OD_600nm_^-1^) was determined using impedance flow cytometry as recently described (Hartmann *et al*., 2021). Arrows indicate the addition of sodium-hypochlorite (NaOCl; 35 mM) Error bars represent standard deviation from at least three replicates.

Accordingly, the experiment was conducted as stated above and immediately after the addition of PBS or the respective sub-lethal doses of NaOCl (35 mM final concentration), samples were taken to assess the oxidative stress level *via* the OxD value of the Mrx1-roGFP2 sensor protein, the DNA-stress level *via* the emission intensity at 640 nm (excitation at 600 nm), and growth via the OD_600nm_. Growth of *C. glutamicum* WT_Mrx1-roGFP2 (pJC1_P*_recA__e2-crimson*) (untreated) proceeded with a growth rate of 0.37 ± 0.01 h^-1^ to a final optical density of 23.1 ± 0.77 (Fig. 3c).

Addition of NaOCl caused an immediate cease of growth but continued growing after one hour at a reduced growth rate of 0.21 ± 0.02 h^-1^ to a final optical density of 22.10 ± 1.43 (Fig. 3c). The transiently impaired growth indicates that the cells experienced oxidative stress and indeed an increased oxidative stress level was detected *via* the redox biosensor protein Mrx1-roGFP2. Mrx1-roGFP2 was more oxidized initially after applying NaOCl to the culture (0.70 - 0.75), when compared to the Mrx1-roGFP2 signals measured for the untreated culture (0.50 - 0.53) (Fig. 3c). Three hours after applying NaOCl, the OxD values decreased to 0.53. This is similar to OxD values determined for non-treated cultures throughout the experiment (Fig. 3c). To compare the DNA-damage response of treated and untreated cultures, the relative fluorescence intensity at 640 nm (normalized to the OD_600nm_) derived from the treated culture was divided by the fluorescence signal intensity determined for the untreated cultures. Consequently, the induction of the DNA-stress reporter plasmid pJC1_P*_recA__e2-crimson* results in values above 1, whereas 1 represents a status where no difference with respect to the induction level between the both approaches was detected. One hour after oxidative stress was applied, the E2-Crimson ratio (treated/untreated) increased from initially (at the time-point of the treatment) 1.11 ± 0.09 to values between 1.3 - 1.5 after five, six and seven hours of cultivation (Fig. 3e). After this, the ratio regenerated back to 1.12 ± 0.05 and was maintained at approximately 0.90 - 1.05 for the rest of the experiment (Fig. 3e).

Growth of the MSH-deficient strain *C. glutamicum* Δ*mshC*_Mrx1-roGFP2 (pJC1_P*_recA__e2-crimson*) proceeded at a growth rate of 0.41 ± 0.06 h^-1^ until the untreated culture reached a final optical density of 23.7 ± 1.5 (Fig. 3d). Similar to *C. glutamicum* WT_Mrx1-roGFP2 (pJC1_P*_recA__e2-crimson*), addition of the artificial oxidant NaOCl resulted in an immediate cease of growth of *C. glutamicum* Δ*mshC*_Mrx1-roGFP2 (pJC1_P*_recA__e2-crimson*). Growth of *C. glutamicum* Δ*mshC*_Mrx1-roGFP2 (pJC1_P*_recA__e2-crimson*) restarted two hours after NaOCl addition and proceeded with a growth rate of 0.18 ± 0.06 h^-1^ until a final optical density of 21.0 ± 1.9 was reached. This is a slightly lower final biomass when compared to the untreated sample (Fig. 3d). In general, the OxD values of the sensor Mrx1-roGFP2 were determined to be highly oxidized for *C. glutamicum* Δ*mshC*_Mrx1-roGFP2 (pJC1_P*_recA__e2-crimson*) (OxD values of 0.90 - 0.99; representing a fully oxidized state of the biosensor). Such high OxD levels were expected for the MSH-deficient strain background, as such values have also been observed in previous studies in absence of any externally applied oxidative stress (Tung *et al*., 2019; Hartmann *et al*., 2020). It was hypothesized, that the absence of MSH leads to an elevation of intracellular ROS levels which in turn induces an auto-oxidation of the biosensor protein Mrx1-roGFP2 (Tung *et al*., 2019; Hartmann *et al*., 2020).This means that the biosensor protein is close to its detection limit due to its almost fully oxidized state and a further induction of oxidative stress cannot adequately be captured by the biosensor. As a result, the OxD values obtained from the NaOCl treated cultures of *C. glutamicum* Δ*mshC*_Mrx1-roGFP2 (pJC1_P*_recA__e2-crimson*) were determined to be fully oxidized at all time-points in this experiment (Fig. 3b). Due to the lack of MSH as the major LMW-thiol in *C. glutamicum* (Reyes *et al*., 2018; Tung *et al*., 2019), an increased susceptibility towards ROS and in turn DNA-damage response was expected for *C. glutamicum* Δ*mshC*_Mrx1-roGFP2 (pJC1_P*_recA__e2-crimson*). Indeed, fluorescence measurements revealed an up to 2.5-fold increase of the E2-Crimson ratio for the NaOCl-treated culture of *C. glutamicum* Δ*mshC*_Mrx1-roGFP2 (pJC1_P*_recA__e2-crimson*) when compared to the signals determined for the control culture. (Fig. 3f). The highest ratio was measured after six and seven hours of cultivation, corresponding to three and four hours after inducing the artificial oxidative stress (Fig. 3d). Afterwards, the ratio of the signal of P*_recA__e2-crimson* returned back to approximately one and remained at this level throughout the experiment (Fig. 3f).

Taken together, the results show that NaOCl-induced oxidative stress indeed coincidences with an increased induction of the DNA-stress reporter P*_recA__e2-crimson*. More pronounced effects of the NaOCl-triggered induction of P*_recA__e2-crimson* were observed in a MSH-deficient mutant strain of *C. glutamicum*, which is known be more susceptible to oxidative stress (Chi *et al*., 2014; Hartmann *et al*., 2020). Consequently, this probably also caused oxidative DNA-damage and a higher loss of the culture’s viability. Thus, the results show that oxidative stress in turn induces DNA stress/damage in *C. glutamicum* and that MSH is a key player for the organisms’ to protect DNA under these conditions.

### 3.5 Effects of the DNA-damaging agent MMS on the redox-potential in *C. glutamicum*

Oxidative stress can cause significant damages to macromolecules such as DNA. Thus, the linkage of oxidative stress and DNA-damage response is well known. In opposite to this, less is known about the link between DNA-damage and oxidative stress. It was demonstrated that exposure to the DNA damaging agent MMS caused a dose-dependent increase in intracellular ROS levels in yeast (Salmon *et al*., 2004; Kitanovic and Wölfl, 2006). Unlike oxidants, MMS is not converted to ROS directly and by this causing oxidative DNA-damage. Thus, this observation indicated that DNA-damage alone can induce an oxidative stress response in yeast (Salmon *et al*., 2004; Kitanovic and Wölfl, 2006). To the best of our knowledge this phenomenon has hitherto not been investigated in *C. glutamicum*.

Prior to studying this in more detail by the use of the double biosensor strains of *C. glutamicum*, the stability of the sensor signal derived from the redox biosensor protein Mrx1-roGFP2 in presence of the DNA damaging agent MMS was tested *in-vitro*. Therefore, crude cell extracts prepared from *E. coli* BL21 (*DE3*) (pET11b_*mrx1-roGFP2*) were used. The addition of MMS to the cell extracts did not alter the biosensor ratio, whereas the addition of reductants or oxidants has led to reduced or increased biosensor ratios, respectively (Fig. 2S). These results indicate that MMS itself does not react with the sensor protein Mrx1-roGFP2. As shown, the presence of the fluorescent protein E2-Crimson did not alter the OxD value of the Mrx1-roGFP2 biosensor protein *in-vitro* (Fig. 1b) as well as *in-vivo* (Table 2). Thus, we concluded that the double sensor set-up is appropriate to analyze potential effects of the DNA-damaging agent MMS on the redox-potential in *C. glutamicum*.

The sensor-equipped strains *C. glutamicum* WT_Mrx1-roGFP2 (pJC1_P*_recA__e2-crimson*) and *C. glutamicum* Δ*mshC*_Mrx1-roGFP2 (pJC1_P*_recA__e2-crimson*) were cultivated in presence of 100 μg/mL and 800 μg/mL MMS. As shown before, these two concentrations were sufficient to adequately induce a DNA-damage response via the reporter plasmid P*_recA__e2-crimson* in both *C. glutamicum* WT_Mrx1-roGFP2 (pJC1_P*_recA__e2-crimson*) and *C. glutamicum* Δ*mshC*_Mrx1-roGFP2 (pJC1_P*_recA__e2-crimson*) (Fig. 2a, b). Final biomass and viability of both strains were only slightly reduced in cultivations with 100 μg/mL MMS when compared to cultivations in absence of any MMS (Table 3). In opposite to this, the population viability was reduced by approximately 50% and 75% in presence of 800 μg/mL MMS for *C. glutamicum* WT_Mrx1-roGFP2 (pJC1_P*_recA__e2-crimson*) and *C. glutamicum* Δ*mshC*_Mrx1-roGFP2 (pJC1_P*_recA__e2-crimson*), respectively (Table 3). The results show that the absence of the antioxidant MSH has led to a higher susceptibility of *C. glutamicum* towards the DNA damaging agent MMS. This implies an interconnection of DNA-damage and oxidative stress in *C. glutamicum*. In order to decipher this interconnection in more detail, the OxD values of the redox biosensor protein in the double biosensor strains were analyzed. In presence of MMS, *C. glutamicum* WT_Mrx1-roGFP2 (pJC1_P*_recA__e2-crimson*) revealed a strong oxidative shift when compared to OxD values determined for cells cultivated in absence of MMS (Fig. 4a). The OxD value in cells of *C. glutamicum* WT_Mrx1-roGFP2 (pJC1_P*_recA__e2-crimson*), cultivated in presence of 800 μg/mL MMS, even reached the highly oxidized states which have been measured for cells lacking MSH (*C. glutamicum* Δ*mshC*_Mrx1-roGFP2 (pJC1_P*_recA__e2-crimson*)). In detail, OxD values were measured to be 0.41 ± 0.01, 0.52 ± 0.01 and 0.86 ± 0.1 for samples without MMS, 100 μg/mL and 800 μg/mL, respectively (Fig. 4a). This corresponds to a redox potential determined for MSH of −285 ± 0.3 (no supplements), −279 ± 0.5 (100 μg/mL) and −253 ± 16 (800 μg/mL) (Fig. 4b).

**Figure 4:**
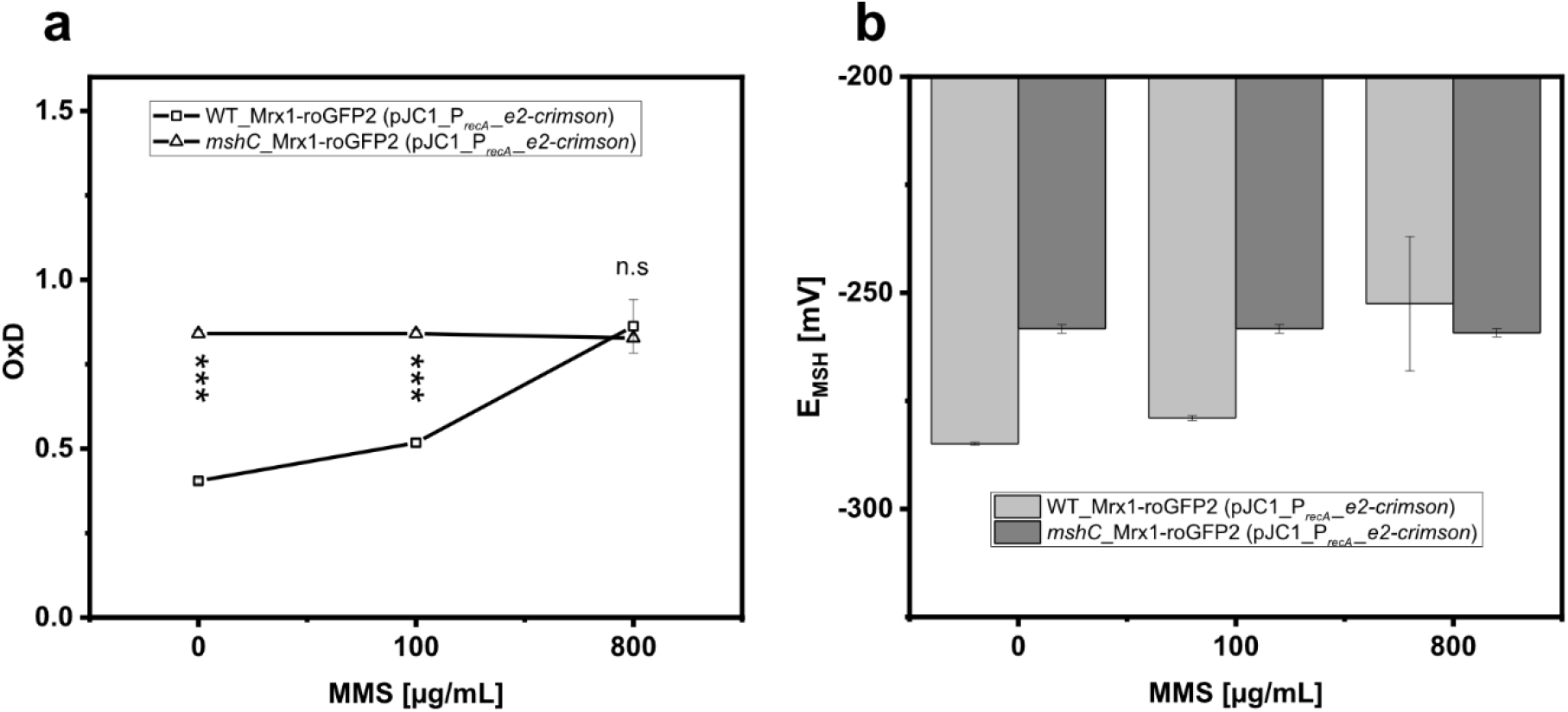
Oxidation degree (OxD) of the redox sensor protein Mrx1-roGFP2 in the double biosensor strain *C. glutamicum* WT_Mrx1-roGFP2 (pJC1_*P_recA__e2-crimson*) and *C. glutamicum* Δ*mshC*_Mrx1-roGFP2 (pJC1_*P_recA__e2-crimson*) **(A)** and corresponding mycothiol redox potentials after cultivation for 24 hours in shaker flasks in presence of different methyl-methanesulfonate (MMS) concentrations **(B)**. Mrx1-roGFP2 fluorescence signal was measured by recording the emission intensity at 510 nm upon excitation at 380 nm and 470 nm. The ratiometric signal was calculated by dividing the former emission intensity by the latter followed by calculating the OxD value and mycothiol redox potential (E_MSH_) according to Equation1 and 2, respectively (Material and Methods). All fluorescence measurements were conducted in black flat-bottomed 96-well microtiter plates using a plate reader device (SpectraMaxiD3). Statistical analysis was performed via One-Way-ANOVA followed by a Tukey’s test (^n.s^ p > 0.05; * p ≤ 0.05; ** p ≤ 0.01; *** p ≤ 0.001).

**Table 3:** Final optical density (OD_600nm_) and viability of *C. glutamicum* WT_Mrx1-roGFP2 (pJC1_P*_recA__e2-crimson*) and *C. glutamicum* Δ*mshC*_Mrx1-roGFP2 (pJC1_P*_recA__e2-crimson*) upon exposure to different methyl-methanesulfonate (MMS) concentrations for 24 hours. Viability was measured via impedance flow cytometry. Viability was calculated (viable cells*OD_600nm_^-1^*mL^-1^) followed by normalizing to the respective viability in absence of any MMS resulting in relative viabilities.

| MMS [μg/mL] | 0 |  | 100 |  | 800 |  |
| --- | --- | --- | --- | --- | --- | --- |
| Strains/ OD<sub>600nm</sub> & Viability | OD<sub>600nm</sub> | Viability [%] | OD<sub>600nm</sub> | Viability [%] | OD<sub>600nm</sub> | Viability [%] |
| WT_Mrx1-roGFP2 (pJC1_P<sub>recA</sub>-e2-crimson) | 5.9 ± 0.3 | 100 | 5.0 ± 0.4 | 85 ± 0 | 3.2 ± 0.1 | 47 ± 9 |
| ΔmshC_Mrx1-roGFP2 (pJC1_P<sub>recA</sub>-e2-crimson) | 5.6 ± 0.2 | 100 | 5.0 ± 0.3 | 83 ± 4 | 3.0 ± 0.2 | 23 ± 16 |

As expected, the MSH-deficient mutant strain Δ*mshC*_Mrx1-roGFP2 (pJC1_P*_recA__e2-crimson*) was almost fully oxidized even without any supplements and OxD levels were maintained between 0.83-0.85, irrespectively of the externally applied MMS concentration (Fig. 4a). This corresponds to redox potentials of approximately −260 mV for all approaches (Fig. 4b). These values are similar to the redox potentials measured for the WT_Mrx1-roGFP2 (pJC1_P*_recA__e2-crimson*) strain cultivated in presence of 800 μg/mL MMS. To note, for mutant strains lacking MSH, the biosensor was almost fully oxidized and thus close to its detection limit. Consequently, an oxidative stress response is difficult to capture. However, when compared to the wild type strain, the cultures viability was reduced in presence of 800 μg/mL MMS indicating a higher susceptibility towards DNA-damage and the concomitant cause of oxidative stress. In agreement to this, an oxidative redox shift in *C. glutamicum* has been observed when facing high doses of MMS.

Taken into account that MMS is not converted to ROS directly and the insensitivity of the redox biosensor protein Mrx1-roGFP2 towards the DNA-damage inducing agent MMS, the observed oxidative shift has to be interconnected to cellular redox changes. The underlying mechanism might be due to an accumulation of ROS upon facing excessive amounts of unrepaired DNA-damage as demonstrated in yeast (Salmon *et al*., 2004; Kitanovic and Wölfl, 2006), however the biochemical causes and mechanism remain to be investigated in future studies.

### 3.6 Effects of the inhibition of cell wall biosynthesis on redox-potential and DNA-damage response in *C. glutamicum*

Several bactericidal antibiotics *e.g*. ampicillin (β-lactam), kanamycin (aminoglycoside), and norfloxacin (fluorchinolon) were recently shown to trigger the oxidative stress response and/or make the cells more susceptible to damages caused by oxidative agents in bacteria such as *E. coli; Acinetobacter baumanii, Rhodococcus equii*, and *Mycobacterium tuberculosis* (Kohanski *et al*., 2007; Bhaskar *et al*., 2014; Dwyer *et al*., 2014; Ajiboye *et al*., 2018; Mourenza *et al*., 2020). These observations underlie the still controversially discussed hypothesis that aforementioned classes of antibiotics in addition to their primary mode of action also cause the generation of lethal ROS (Dwyer *et al*., 2015; Imlay, 2015; Van Acker and Coenye, 2017). Sensor based-analyses of oxidative stress *e.g. via* roGFP based sensors in combination with a second sensor *e.g*. for DNA-damage might help to decipher the complexity of antibiotic lethality. Addition of sub-lethal doses of the β-lactam antibiotic penicillin causes growth inhibition in *C. glutamicum* and triggers glutamate production (Nara *et al*., 1964; Eggeling *et al*., 2001). Using the sensor equipped strains *C. glutamicum* WT_Mrx1-roGFP2 (pJC1_P*_recA__e2-crimson*) and *C. glutamicum* Δ*mshC*_Mrx1-roGFP2 (pJC1_P*_recA__e2-crimson*), we aimed to investigate for possible effects of penicillin addition on both oxidative stress and the DNA-damage response in *C. glutamicum*.

For this purpose the two sensor-equipped strains were cultivated in presence of 0.1, 0.5, 4, and 16 U/mL penicillin, as at these concentrations growth of *C. glutamicum* was inhibited but the majority of the cells maintain viable for glutamate production (Hartmann *et al*., 2021). For all tested penicillin concentrations, the final OD_600nm_ of *C. glutamicum* WT_Mrx1-roGFP2 (pJC1_P*_recA__e2-crimson*) was reduced by around 50% when compared to cultivations conducted in absence of penicillin (Fig. 5a). For *C. glutamicum* Δ*mshC*_Mrx1-roGFP2 (pJC1_P*_recA__e2-crimson*), the final OD_600nm_ was reduced by approximately 50% in presence of 0.1 U/mL and 0.5 U/mL. When compared to the wild type strain, a further increase of the penicillin concentration in turn resulted in a significant reduction of the final OD_600nm_ (Fig. 5a). To measure the oxidative stress “status” of the cultures, the oxidation state of the redox biosensor protein Mrx1-roGFP2 was determined. The OxD values in cells of *C. glutamicum* WT_Mrx1-roGFP2 (pJC1_P*_recA__e2-crimson*) were already increased from 0.34 ± 0.10 (in absence of penicillin) to 0.80 ± 0.17 in presence of 0.1 U/mL penicillin (Fig. 5b), indicating the utilization of MSH as antioxidant under these conditions. For the MSH-deficient strain *C. glutamicum* Δ*mshC*_Mrx1-roGFP2 (pJC1_P*_recA__e2-crimson*), an increase from an OxD value of 0.63 ± 0.00 (in absence of penicillin) to a fully oxidized state of the redox biosensor protein Mrx1-roGFP2 (0.99) was observed in presence of 0.1 U/mL penicillin (Fig. 5b). Consequently, exposure to sub-lethal concentrations of penicillin resulted in an oxidative redox shift (according to Equation 2; Nernst equation) in both *C. glutamicum* WT_Mrx1-roGFP2 (pJC1_P*_recA__e2-crimson*) and *C. glutamicum* Δ*mshC*_Mrx1-roGFP2 (pJC1_P*_recA__e2-crimson*) (Fig. 5c). To note, crude cell extracts prepared from *E. coli* BL21 (*DE3*) (pET11b_*mrx1-roGFP2*) were used to test whether penicillin itself affects the oxidation state of the protein. However, the addition of penicillin to the cell extracts did not alter the biosensor ratio, whereas the addition of reductants or oxidants has led to reduced or increased biosensor ratios, respectively (Fig. 2S). This indicates that MSH as non-enzymatic antioxidant and major LMW-thiol in *C. glutamicum* might play a role in protecting the cell against oxidative stress when becoming exposed to the antibiotic penicillin.

**Figure 5:**
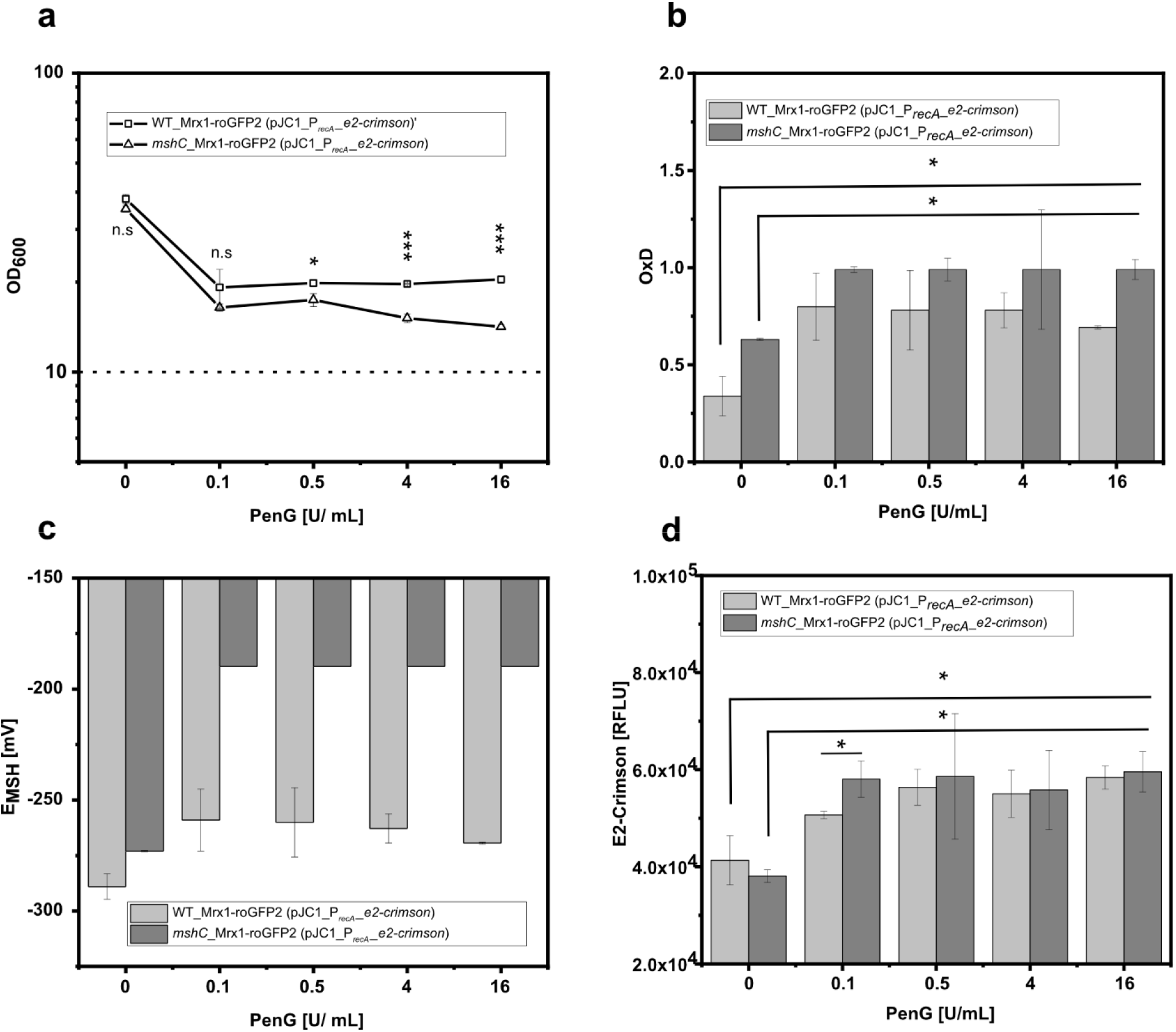
Exposure of the double biosensor strains *C. glutamicum* WT_Mrx1-roGFP2 (pJC1_P*_recA__e2-crimson*) and *C. glutamicum* Δ*mshC*_Mrx1-roGFP2 (pJC1_P*_recA__e2-crimson*) to different PenicillinG (PenG) concentrations. Final optical density (OD_600nm_) **(A)**, oxidation degree (OxD) of the redox biosensor protein Mrx1-roGFP2 **(B)**, the respective mycothiol redox potential (EMSH) **(C)**, and the E2-Crimson fluorescence intensities **(D)**. Dashed line indicates the initial set OD_600nm_. Growth experiment was conducted in CGXII minimal medium (2% glucose) in FlowerPlates. Mrx1-roGFP2 fluorescence signal was measured by recording the emission intensity at 510 nm upon excitation at 380 nm and 470 nm. The ratiometric signal was calculated by dividing the former emission intensity by the latter followed by calculating the OxD value according to equation1 (Material and Methods). E2-Crimson fluorescence intensity was recorded at 640 nm upon an excitation at 600 nm. All fluorescence measurements were conducted in black flat-bottomed 96-well microtiter plates using a plate reader device (SpectraMaxiD3). Error bars represent standard deviation from at least three replicates. Statistical analysis was performed via One-Way-ANOVA followed by a Tukey’s test (^ns^ p > 0.05; * p ≤ 0.05; ** p ≤ 0.01; *** p ≤ 0.001).

To further test the DNA-stress “status”, the reporter activity of the DNA-damage biosensor P*_recA__e2-crimson* was measured in both *C. glutamicum* WT_Mrx1-roGFP2 (pJC1_P*_recA__e2-crimson*) and *C. glutamicum* Δ*mshC*_Mrx1-roGFP2 (pJC1_P*_recA__e2-crimson*). For all tested penicillin concentrations, the reporter activity was significantly increased in both strain backgrounds when compared to cultivations conducted in absence of penicillin (Fig. 5d). At low penicillin concentrations (0.1 U/mL), significantly higher E2-Crimson fluorescence intensities were measured in the MSH-deficient strain *C. glutamicum* Δ*mshC*_Mrx1-roGFP2 (pJC1_P*_recA__e2-crimson*). This indicates that penicillin addition to *C. glutamicum* cultures causes an effect on its non-enzymatic antioxidant defense system. This might be the result of ROS species which have been internally formed in response to the inhibition of growth by the antibiotic.

Taken together, these experiments illustrate how the double sensor system targeting two different types of stress in one cell can be utilized as a powerful analytical tool to decipher the complex interplay in microbial stress responses.

## 4 Conclusions

In this study, we demonstrated the applicability of a combinatorial biosensor set-up based on a redox sensitive fluorescent protein (Mrx1-roGFP2) and a DNA-stress reporter plasmid (pJC1_P*_recA__e2-crimson*) towards assessing oxidative DNA-damage in *C. glutamicum*. Utilizing this combinatorial biosensor concept in a MSH-deficient mutant strain of *C. glutamicum* allowed us to identify the major LMW-thiol and non-enzymatic antioxidant MSH as key player to protect the cells against DNA-damage when facing oxidative stress. Further physiological studies based on this combinatorial biosensor concept revealed that the DNA-damaging agent MMS has an impact on the cellular redox environment. This suggests a direct link of DNA-damage and oxidative stress response in *C. glutamicum*. However, the underlying biochemical mechanism remains unclear and has to be addressed in future studies. Finally, we observed that inhibition of cell wall biosynthesis by sub-lethal doses of penicillin caused an oxidative redox shift and the induction of the DNA-damage response in *C. glutamicum*. This supports the still controversially discussed hypothesis that antibiotics in addition to their primary mode of action also induce oxidative stress and DNA-damage.

The compatibility of the redox biosensor protein Mrx1-roGFP2 with the fluorescent protein E2-Crimson shown here provides the basis to further combine other redox fluorescent reporter proteins such as Grx1-roGFP2 (*E. coli, Saccharomyces cerevisiae;* glutathione) or Brx1-roGFP2 (*Bacillus subtilis*, bacillithiol) with reporter systems based on E2-Crimson. This will facilitate in-depth studies addressing redox-related stress-physiology for a broad portfolio of other relevant platform organisms to support metabolic engineering and bioprocess development in the future.

## Supporting information

SupplData_OXStress_DNAdamage_Cglutamicum

## Credit author statement

**Fabian Stefan Franz Hartmann:** Conceptualization; Data curation, Formal analysis, Investigation, Methodology, Supervision, Validation, Visualization, Writing-original draft; Writing-review & editing; **Ioannis Anastasiou:** Data curation; Formal analysis; Investigation; Validation; Writing-review & editing; **Tamara Weiß:** Data curation; Formal analysis; Writing-review & editing; **Tsenguunmaa Lkhaasuren:** Formal analysis, Writing-review & editing; **Gerd Michael Seibold:** Conceptualization; Funding acquisition; Supervision; Validation; Writing-original draft; Writing-review & editing

## Availability of data and materials

All data generated and analyzed during this study are included in this article and its additional files. Raw datasets are available from the corresponding author on reasonable request.

## Acknowledgment

We would like to thank the Fermentation Core at DTU Bioengineering for excellent technical support and Prof. Dr. Julia Frunzke (Institute of Bio- and Geosciences - IBG-1: Biotechnology, Forschungszentrum Jülich GmbH) for providing the plasmid pJC1_P*_recA__e2-crimson*.

