## Supplementary material for "Combined sensor-based monitoring of mycothiol redox potential and DNA-damage response in *Corynebacterium glutamicum*": SupplData_OXStress_DNAdamage_Cglutamicum

The supplementary material comprises two figures:

**Figure S1:** On-line monitoring of fluorescence (Exc. 610 nm/Em. 640 nm) derived from *C. glutamicum* WT (pJC1_$P_{recA}$-*e2-crimson*) upon applying methyl-methansulfonate (MMS) to the culture.

**Figure S2:** Determination of the redox biosensor protein Mrx1-roGFP2 in vitro upon exposure to different chemicals

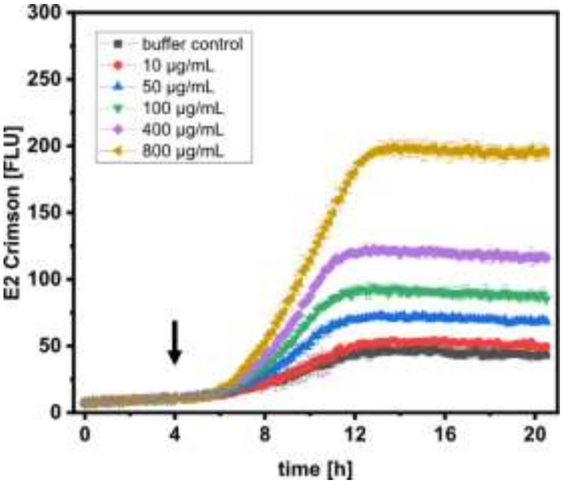


FigureS1: On-line monitoring of fluorescence (Exc. 600 nm/Em. 640 nm) derived from *C. glutamicum* WT (pJC1_$P_{recA}$-*e2-crimson*) upon applying methyl-methansulfonate (MMS) to the culture. Error bars represent standard deviation derived from at least three replicates.

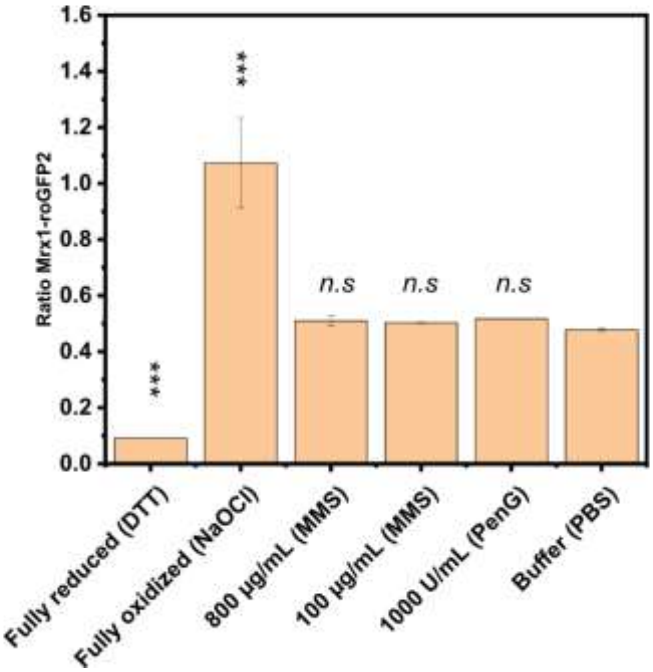


FigureS2: Determination of the redox biosensor protein Mrx1-roGFP2 in vitro upon exposure to different chemicals. Methyl-methanesulfonate (MMS), dithiothreitol (DTT), sodium-hypochlorite (NaOCl), penicillinG (PenG), potassium phosphate buffer (PBS) was applied to cell extracts of *E. coli* (pET11b_*mrx1-roGFP2*) in black-flat bottomed 96-well plates. For reduced and oxidized controls, DTT or NaOCl were applied, respectively. Fluorescence analysis was performed in a microplate reader device (SpectraMax iD3). Mrx1-roGFP2 fluorescence signal was measured by recording the emission intensity at 510 nm upon excitation at 380 nm and 470 nm. The ratiometric signal was calculated by dividing the former emission intensity by the latter. Error bars represent standard deviation derived from at least three replicates. Statistical analysis was performed via One-Way-ANOVA followed by a Tukey´s test ($^{n.s}$ $p > 0.05$; * $p \leq 0.05$; ** $p \leq 0.01$; *** $p \leq 0.001$). Analysis shows significance in difference of treated samples compared to the PBS control.
